# A Predictive Model for Coupling Cell Division Orientation to Tissue Mechanics During Epithelial Morphogenesis

**DOI:** 10.64898/2026.04.17.719304

**Authors:** Somiealo Azote epse Hassikpezi, Rajendra Singh Negi, No (Arnold) Chen, M. Lisa Manning

**Author notes:** These authors contributed equally and share first authorship (Somiealo Azote epse Hassikpezi, Rajendra Singh Negi).

## Abstract

Stratified epithelial tissues such as the skin epidermis maintain barrier integrity during development and homeostasis through the coordinated action of cell proliferation, differentiation, delamination, and tissue-scale mechanical forces. During development, the orientation of cell division within the basal layer plays a pivotal role in tissue stratification; however, the mechanical principles linking the orientation of the division plane to these processes across developmental stages remain poorly understood. Here, we expand a recently developed three-dimensional vertex model for stratified epithelia, composed of the basement membrane, basal, and suprabasal layers, to study the mechanical and structural impact of cell divisions with a wider range of orientations. The model integrates developmental stage via specific changes in heterotypic interfacial tensions (arising from actomyosin cortical contractility and adhesion molecules at the basal–suprabasal interface) and tissue stiffness that have been quantified previously in experiments. By systematically varying background mechanical parameters, we investigate how heterotypic tension, division orientation, and tissue fluidity collectively influence the outcome of cell division. Our goal is to uncover the strategies that the embryo may employ to generate stratified phenotypes at different developmental stages, recognizing that these strategies might evolve over time. Although our focus is on the embryonic developmental stages of the epidermis, this framework may also be extended to investigate transformed cells, such as in cancer, to explore how altered division orientation contributes to precancerous or transformed phenotypes.

## 1 Introduction

Stratified epithelial tissues, such as the skin epidermis, achieve and maintain their barrier function through the coordinated regulation of cell proliferation, differentiation, delamination, and tissue-scale mechanical forces [5, 9, 29]. A defining feature of epidermal development is the orientation of cell division within the basal progenitor layer, which influences both the spatial arrangement of daughter cells and their subsequent fate decisions [14]. Perturbations in division orientation have been implicated in a wide range of pathological conditions, including impaired barrier formation, aberrant stratification, and cancer, underscoring the importance of understanding how division orientation is regulated and how it feeds back onto tissue organization and mechanics[ [30, 3, 13, 33, 46].

Previous genetic and imaging studies established that oriented cell divisions play a central role in epidermal morphogenesis. During embryonic development, basal progenitors often divide perpendicularly or obliquely relative to the basement membrane, producing suprabasal daughters that directly contribute to stratification and differentiation [22, 3]. In contrast, planar divisions expand the basal progenitor pool and support tissue growth. Maintaining the correct balance between these division modes is essential: excessive perpendicular (asymmetric division where orientation biases daughter cells are displaced suprabasally) divisions can prematurely deplete progenitors, whereas excessive planar (symmetric division where orientation biases daughter cells remain basal) divisions impair stratification and barrier formation [30, 46, 22]

In addition, division orientation is correlated with the overall rate of basal cell division, acting in close coordination with fate decisions and tissue mechanics to quantitatively constrain epidermal stratification dynamics. Long-term intravital live imaging of the mouse interfollicular epidermis has shown that under adult homeostatic conditions, basal stem and progenitor cells divide relatively slowly, with a division probability of approximately 0.3 per basal cell per day, corresponding to roughly one division every 2–3 days per cell [34, 27]. These studies further revealed that basal cell divisions are temporally and spatially coupled to differentiation events, such that differentiation of one basal cell is typically followed by a compensatory division of a neighboring basal cell within ∼ 24–48 hours, ensuring robust maintenance of basal cell density and tissue turnover [34, 27]. In mice, this coordinated behavior supports complete epidermal renewal on a timescale of ∼ 8–10 days. During embryonic development, however, basal and suprabasal keratinocytes divide at substantially higher rates, estimated to be about 0.7 divisions per basal cell per day, coinciding with rapid tissue growth and stratification [22, 8]. Under these high-proliferation conditions, stratification can proceed efficiently through a combination of perpendicular divisions and crowding-induced delamination, even when spindle orientation cues are weakened [8]. In contrast, as division rates decline and mechanical resistance in the basal layer increases during later developmental stages, excessive planar divisions fail to generate sufficient suprabasal cells, while excessive perpendicular divisions risk progenitor depletion [30, 46].

At the molecular level, epithelial division orientation is tightly governed by polarity regulators such as atypical protein kinase C lambda (aPKC*λ*). Conditional loss of aPKC*λ* in mouse epidermis disrupts spindle alignment, biases divisions toward perpendicular orientations, and induces progressive defects in stem cell maintenance, particularly within the hair follicle bulge niche [30]. These findings highlight how misregulation of division orientation can drive inappropriate fate choices, ultimately compromising long-term tissue integrity. Across epithelial tissues more broadly, spindle misorientation has been associated with disorganized architecture, abnormal differentiation, and tumorigenesis [33].

Misoriented divisions are not only restricted to developmental defects, but can also be a hallmark of disease. In multiple epithelial cancers, including breast cancer and basal cell carcinoma (BCC), altered division orientation correlates with disrupted tissue architecture, loss of polarity, and invasive behavior [33, 42]. In the context of skin cancer, oncogenic mutations perturb basal cell differentiation programs and division orientation, leading to uncontrolled proliferation within the basal layer. For example, perturbations in organelle inheritance and spindle positioning alter the balance between growth and differentiation in epidermal progenitors [1], while changes in tissue-scale mechanics reshape tumor architecture and functional heterogeneity [13]. Together, these studies underscore the tight coupling between division orientation, fate regulation, and mechanical context in both normal and diseased epithelia.

Although many of the molecular systems that orient the spindle-polarity complexes – the cortical force generator, centrosomal pathways and endocytic regulator – have been well described [10, 4, 19, 18], the mechanical consequences of division orientation remain less well understood. Classical and contemporary reviews have emphasized how division orientation is established even in densely packed epithelia such as the epidermis[46, 10]. However, whether division orientation actively feeds back to regulate tissue mechanics – by relaxing anisotropic stresses, reorganizing force transmission, or stabilizing tissue architecture, rather than merely responding to pre-existing mechanical cues – remains an open question.

Recent experimental work has begun to challenge the idea that division orientation alone dictates stratification. Quantitative lineage tracing and live imaging have shown that epidermal stratification can proceed through high proliferation rates and delamination, even when spindle orientation is perturbed [8, 9, 38]. Similarly, stem cell fate commitment has been shown to occur gradually and can be uncoupled from immediate cell-cycle exit or strict asymmetric division rules [27, 6]. These findings point to a more nuanced, population-level regulation of epidermal homeostasis, in which division orientation, tissue crowding, and mechanical forces jointly shape outcomes.

Recent studies motivate the hypothesis that epidermal stratification is governed by a mechanical feedback loop that links the mechanics of the basal layer and the orientation of the division. In the forward direction along that loop, there is evidence that mechanics help to direct the orientation of division. Tissue geometry and cell shape bias spindle orientation through physical constraints, often aligning divisions with the long axis of the cell [28, 3]. External forces can directly reposition the mitotic spindle, demonstrating that division orientation is mechanically sensitive rather than purely genetically programmed [12].

To predict how these cell/tissue geometries and cell-scale forces arise, recent work has successfully compared experimental observations to three-dimensional biophysical models for epithelial tissues, called vertex or Voronoi models [48, 26, 35, 31, 16, 21]. Applying these models to the developing mouse epithelia, some of us [45] have demonstrated that in early embryonic development (E14.5), the basal layer is fluid-like, mechanical resistance to deformation is low, and the barrier between basal and suprabasal layers is weak. Under these conditions, we speculate that spindle orientation is permissive, allowing frequent oblique and perpendicular divisions as well as facile delamination, resulting in rapid stratification [45, 29, 3, 46, 27, 22]. At later developmental stages (E15.5 and above), the basal layer stiffens, a mechanical barrier emerges at the basal–suprabasal interface, and resistance to deformation and delamination increases[45]. This barrier is characterized by integrin-mediated wetting tensions at the basal–basement membrane interface, heterotypic interfacial tensions between basal and suprabasal cells, and increased basal tissue stiffness. We speculate that as a consequence, division orientation becomes mechanically biased, perpendicular divisions are suppressed, and stratification slows as epidermal homeostasis is achieved [22, 3]. Taken together, this previous work suggests that mechanics may play a role in dictating the orientation of cell divisions in the stratified mouse epithelium.

In addition, mechanics also plays a role in division-independent delamination. It has been shown such the rate of division-independent delamination increases at later stages, as the basal layer and underlying basement membrane stiffen [29, 45]. While some modeling work has suggested that delamination could be directly driven by a mechanical instability as the tissue becomes more crowded [16], recent work by some of us [45] indicates that at later stages (E15.5, E16.5), delamination is instead governed by notch-triggered cell fate decisions. Therefore, an additional open question is how cell divisions impact tissue mechanics, both in cases of increasing cell density and in cases where cell-fate-based delaminations maintain the tissue at constant cell density.

While there is much work in this forward direction – i.e. studying how tissue mechanics impacts division and delamination – our goal here is to ‘close the mechanical feedback loop’ by investigating how cell division impacts tissue geometry, stratification and mechanics. To do so, we extend a three-dimensional vertex model for stratified epithelia to incorporate basal stem-cell divisions. We explicitly prescribe the division angle and systematically vary it from symmetric to asymmetric orientations. This parameter sweep was performed across multiple developmental stages and division rates. Through this approach, we quantify how division orientation shapes key tissue-level properties, including stratification dynamics, mechanical organization, and basal-layer fluidization. Our results provide a quantitative mechanical description of the experimental (and intuitive) observation that division orientation governs the rate at which daughter cells stratify. It also indicates that overall tissue mechanics are affected by rapid cell divisions, and that both these effects are modulated by developmental changes in surrounding tissue mechanics. Together, this work establishes a mechanistic framework linking division orientation to tissue mechanics, providing new insight into how embryos regulate epidermal stratification during development and how disruptions of this coupling may contribute to disease, including cancer.

## 2 Methods

### 2.1 Simulation Methods

We implement a 3D vertex model in which cells are polyhedral objects with prescribed homeostatic volume *V*_0_ and surface area *S*_0_ [48, 47, 26]. We use the open source code developed by Zhang and Schwarz as a starting point for our extended model [48]. The tissue dynamics arise from equations of motion applied to the degrees of freedom, i.e. the vertices. The epithelial model contains two key compartments: a suprabasal differentiated layer and a one-cell thick basal stem cell layer. As the vertex framework uses space-filling (confluent) polyhedra, we represented the basement membrane as an additional interconnected layer positioned just below the basal cells. The energy functional of such shaped-based interactions is given by:

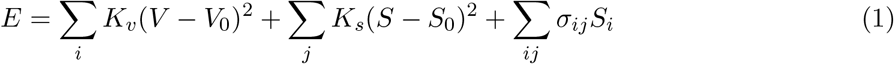

Here, the first term of the energy functional represents the mechanical resistance of cells to volumetric perturbations. The second term captures the competition between actomyosin contractility, which drives cells toward smaller surface areas, and cadherin-mediated adhesion, which promotes the lengthening of cell-cell contacts. The third term represents the heterotypic interfacial tensions that arise between two different cell types [41, 36]. In our system, this term is used to represent the interfacial tension between basal and suprabasal cells and the integrin-driven tension that regulates wetting behavior at the basal-basement membrane interface. The parameters *K*_*V*_ and *K*_*S*_ denote the elastic moduli associated with the volume and surface area of the cells, respectively. Upon non-dimensionalization, this energy functional allows the definition of a key order parameter known as the target shape index, 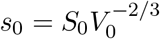 [26], which controls the mechanical rigidity of the tissue.

The system evolves under overdamped Langevin dynamics, with vertex positions continuously updated to reduce the total mechanical energy.

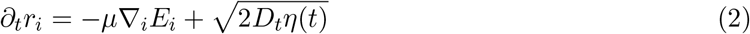

The term on the left-hand side represents the viscous drag forces. On the right-hand side, the first term describes the interaction forces arising from the energy potential times a mobility or inverse drag *μ*, while the second accounts for white noise fluctuations *η*(*t*) in the vertex positions *r*_*i*_, of strength *D*_*t*_. Such fluctuations are a simple, memory-less approximation to cell-generated active forces, such as active tensions and cell motility forces, which are observed in experiments. See SI Section 1 and Table S1 for a list of model parameter used in the simulations.

In simulations where cell divisions are implemented, at each division interval *t*_*div*_, a growing basal cell (defined as a basal cell with a non-zero growth coefficient) is selected to undergo division. We study division times ranging from *t*_*div*_ = 50 (fastest rate of division) to 800 (slowest rate of division). This ensures that at most one cell divides during a single time step, thereby preventing simultaneous division events. We typically simulate about 150 cells in the basal layer, so the shortest timescale *t*_*div*_ = 50 corresponds to a rate of division of 1*/*(50*τ*) per 150 cells. As highlighted in the results section below, *τ* ≈ 0.1 min, corresponding to a rate of ≈ 0.001 divisions per basal cell per minute, or about two divisions per basal cell per day.

As highlighted in the introduction, basal cell division rates in mouse epithelial have been measured in experiments ranging from fast rates of about 0.7 divisions per cell per day during developmental stages E14-E16 [22, 9] to about 0.3 divisions per basal cell per day during homeostasis in adults [27]. These correspond to *t*_*div*_ values of about 150 and 300, respectively. Therefore, the range of cell division rates we study is in the biophysically relevant regime, and *t*_*div*_ between 100 and 200 is likely the relevant division timescale for the E14-E16 developmental stages we study. We express cell division in terms of the basal-cell division rate *λ*. In simulation units, *λ* is measured in units of 1*/τ* per cell and, for a basal layer containing *N*_basal_ ≈ 150 cells, is related to the division interval *t*_div_ by

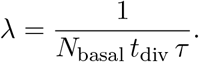

Using *τ* ≈ 0.1 min, this corresponds to a physical division rate

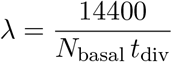

in units of divisions per basal cell per estimated day. Thus, unless stated otherwise, *λ* is given in units of divisions per basal cell per estimated day (1 day = 14400 *τ*).

The division plane is defined by two angles *θ, ϕ*, which specify the orientation of the mitotic axis. The angle *θ* is measured with respect to basal layer: *θ* = 0 corresponds to an in-plane (symmetric) division, whereas *θ* = *π/*2 corresponds to a perpendicular (asymmetric division). The angle *ϕ* ∈ [0, 2*π*] is the azimuthal angle about the basal plane normal and rotationally symmetric, we set *ϕ* = 0 without loss of generality.

During a division event, the parent cell is replaced by two daughter Voronoi seeds that are displaced along the specified mitotic axis. The full Voronoi tessellation is then recomputed, and the entire tissue topology is reconstructed accordingly. Physical properties are assigned to the daughter cells such that volume is conserved, and both cells initially inherit the basal identity of the parent cell. Then the vertex energy (Eq. 1) is re-minimized before normal dynamics according to Eq. 2 are resumed.

This procedure ensures that cell division is implemented in a geometrically exact and topologically consistent manner while remaining fully compatible with the vertex-based mechanical model of the tissue. To maintain continuous tissue turnover, we enforce that at least one basal cell is actively growing at all times. At prescribed intervals, a basal cell is randomly selected to receive a non-zero volume-growth coefficient, ensuring that division events occur intermittently rather than synchronizing across the tissue. If the designated growing cell stratifies before division, a new basal cell is immediately selected to replace it, thereby sustaining basal proliferation.

### 2.2 Analysis methods

We also wish to quantify how cells move and change neighbors during our simulation, especially after a cell division event. Although the mean squared displacement is a standard metric to quantify behavior of cell motion, it includes global tissue motions that are not relevant for neighbor exchange and averages over many time windows. To filter out these global tissue motions and identify the cell division time as a key starting point, we instead measure the relative squared separation between daughter cells, which quantifies how cell division drives rearrangements in the basal layer following a division event. Upon division at time *t*_0_, we record the initial positions, and trajectories of the two daughter cells, *d*_1_ and *d*_2_, identified with their centroid positions 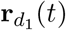 and 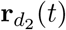 and then define the instantaneous separation vector as

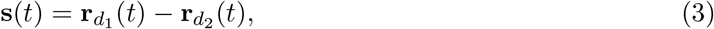

and its value at the time of division

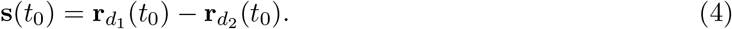

We next track the change in the separation vector

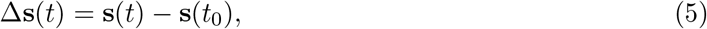

which measures how the daughter pair deforms, rotates, and relaxes relative to its initial configuration.

From this, we define the daughter-daughter separation squared

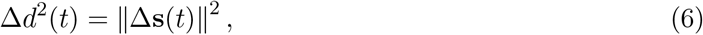

where the average ∥·∥ is taken over an ensemble of all cell division events. In contrast to a traditional mean-squared displacement calculation which is time-invariant, there is no average taken over time. The quantity *t* is the time since the division event.

## 3 Results

### 3.1 Constraining the natural timescale for 3D vertex models

In order to estimate how natural time units in our simulations compare to actual time units in experiments, we turn to experiments that measure rheology using ferrofluidic droplets. As these droplets have not been deployed in mouse embryos, we focus instead on rheological data from zebrafish embryos[40]. In these experiments, a single ferrofluid droplet is placed between cells in the progenitor zone or presomitic mesoderm of developing zebrafish embryos. When the droplet is actuated, it creates a local force dipole that deforms nearby tissue and does not cause visible cell rearrangements. By analyzing how the tissue responds over time, the authors were able to identify two main relaxation timescales: 1s and 1min.

Using Eq. 1, we model a droplet in a confluent tissue of (*n* = 2115) cells by representing the droplet with vertex cells (*n* = 17). We keep the target shape index of the droplet Vertex cells fixed in the fluid-like regime, *s*_droplet_ = 5.8, where they do not contribute to the shear modulus because the effective interfacial tension between cell interfaces is zero in this regime [26, 48, 21]. To distinguish the droplet cells from the surrounding cells, we introduce an explicit hetrotypical interfacial tension *σ* as defined in Eq 1. The droplet is deformed without any net force, and its motion is governed by the equation of motion [11]

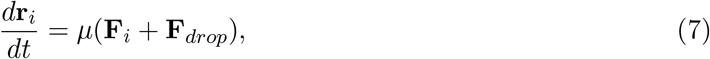

where

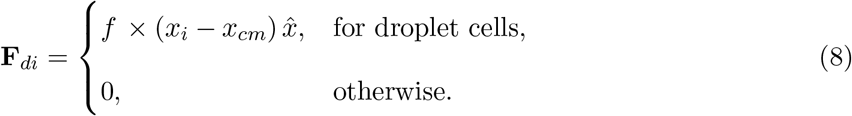

and **F**_*i*_ = ∇_*i*_*E* is the gradient of energy functional. Here, *f* is proportional to the constant applied force that deforms the droplet along the 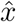 direction, and *x*_*cm*_ denotes the center of mass of the droplet. When the force *f* is applied, the droplet relaxes into an ellipsoidal shape aligned with the direction of *f*. After the force is removed, the droplet returns to its original spherical shape. We quantify the deformation using the strain 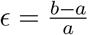, where *a* and *b* are the major and minor axes of the ellipsoid, respectively.

The strain relaxation as a function of time during a single cycle, shown in Figure 1, exhibits at least two distinct characteristic timescales. Such bimodal relaxation behavior is commonly modeled using combinations of springs and dashpots. In this case, the response is best described by a generalized Maxwell fluid consisting of a single spring in parallel with two dashpot elements [40, 15].

**Figure 1:**
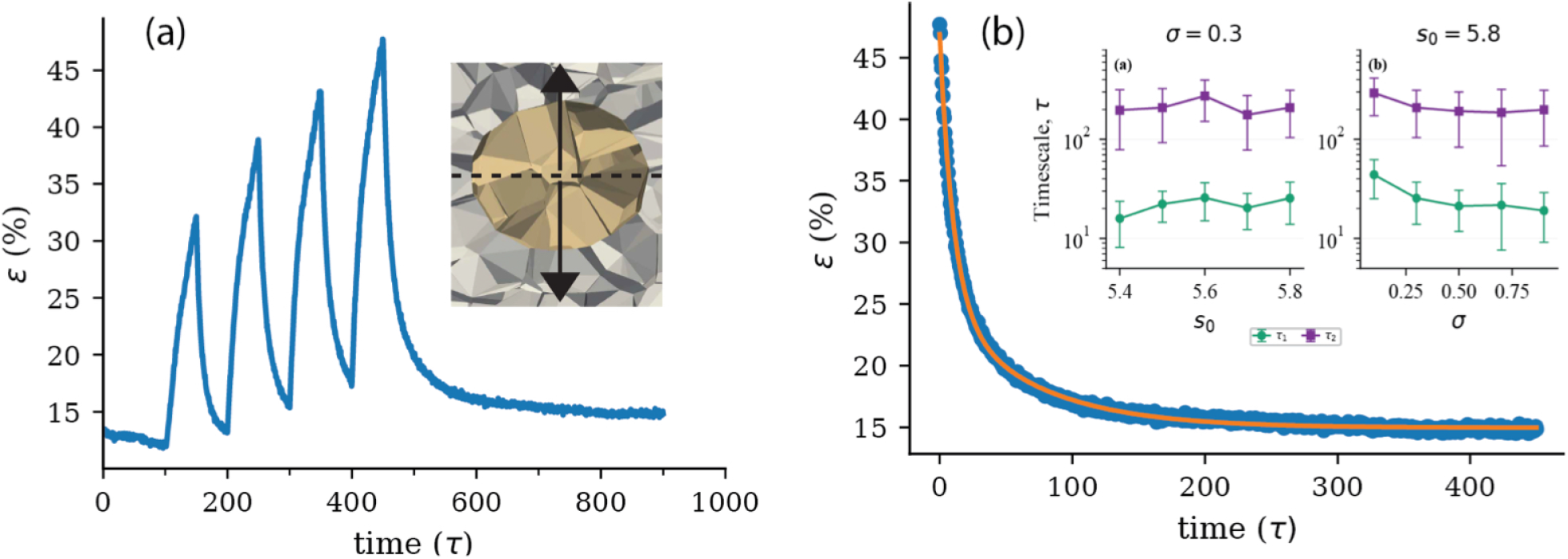
(a) Strain evolution over a series of actuation cycles.The droplet relaxes back to its initial shape after the force is turned off. The inset of (a) is a snapshot of the geometry of the simulations. The dashed line denotes the midsection of the droplet, and arrows represents the direction of the deformation. (b)An example strain–relaxation curve (circles) is fitted to a rheological model (solid line) exhibiting two characteristic relaxation timescales,*τ*_1_ = 10.0*τ* and *τ*_2_ = 68*τ*, for parameters *s*_0_ = 5.8, *σ*_*ij*_ = 0.80, and *f* = 0.50.The inset in panel shows the relaxation timescales as a function of the target shape index *s*_0_ and the droplet surface tension *σ*_*ij*_, green squared denote the shorter timescale *τ*_1_, while magenta squares denote the longer timescale *τ*_2_. Error bars are obtained from 40 independent simulation runs.

We observe that the longer timescale is not strongly dependent on either the tissue stiffess *s*_0_ or the the droplet surface tension *σ* (Fig 1(b) inset). This indicates that 3D vertex models have a natural time unit that does not vary strongly with tissue mechanics, suggesting that numbers extracted from comparison to zebrafish embryos may indeed be relevant for mouse embryos. The longer relaxation timescale extracted from the simulations is compared with the longer timescale measured in zebrafish tailbud tissue, which is approximately 1 min. This suggests a natural simulation time unit of *τ* = 0.1min or 10*τ* = 1min within an order of magnitude.

In previous work, this timescale was independently estimated by computing the velocity of Kupffer’s vesicle (KV), the left–right organizer in zebrafish [37]. The characteristic speed of KV is approximately 0.55 *μ*m*/*min. Using a typical cell size of *l* ∼ 10 *μ*m, this corresponds to a physical speed of ∼ 0.055 *l/*min. In the 3D vertex model, with a characteristic timescale *τ* = 0.1 min, this yields a dimensionless speed *v*_KV_ ∼ 0.0055 *l/τ*, which is consistent with values obtained from vertex-based simulations [24]. Similarly, in Voronoi model simulations, where the natural unit of time is on the order of ∼ 1 min, the corresponding dimensionless speed is *v*_KV_ ∼ 0.055 *l/τ*. This estimate is in agreement with speeds reported in Voronoi-based simulations [37, 11]. In summary, while the inferred natural timescales differ between the vertex (*τ* = 0.1min) and Voronoi (*τ* = 1min) frameworks, the predicted physical speeds are in agreement with each other and with experimental measurements, adding additional support to both estimates.

### 3.2 Developmental stage-specific epidermal tissue mechanical parameters

Recently, some of us quantitatively inferred the stage-dependent mechanical barrier between the basal and suprabasal layers of the stratified mouse epidermis. We combined quantitative analysis of cell and tissue shapes from experimental imaging with those same quantities in large parameter sweeps in 3D vertex model simulations, in order to pinpoint how key model parameters change across developmental stages (E14-E16) [45]. The important mechanical parameters governing tissue mechanics are the heterotypic interfacial tension *σ*_*a*_, the basal-basement membrane wetting tension *σ*_*b*_, and the basal layer stiffness Δ*s*. These parameters collectively define a quantitative mechanical barrier at the basal-suprabasal interface, which emerges as a pivotal cellular mechanism regulating cell rearrangements and delamination during epidermal multilayering and renewal. The strength of this barrier increases from E14 to E16 as the basal stem cell layer transitions from a fluid-like to a progressively stiffer state. In addition, there is an increase in heterotypic interfacial tension at the basal-suprabasal interface, associated with differential adhesion (including actomyosin cortical tension and E-cadherin–mediated adhesion), while the wetting mediated by integrin adhesion at the basal–basement membrane interface remains similar between E14 and E16.

The previously quantified, stage-specific mechanical barrier defines a calibrated mechanical landscape for studying stratification dynamics. In this work, we fix the experimentally inferred parameters (*σ*_*a*_, *σ*_*b*_, Δ*s*) at two developmental stages (E14: (0.044, 0.062, 0); E16: (0.116, 0.067, 0.14)) within the 3D vertex model. Because the tissue at E15 is close to a mechanical droplet-forming instability that can be nucleated by cell division [45], we avoid that complication here and focus on differences between E14 and E16.

Before introducing cell division, we first test whether the mechanical barrier between basal and suprabasal layers is sufficient to orient the division plane. This allows us to isolate the role of the mechanical environment in controlling basal stem cell division orientation.

### 3.3 Geometry under growth: Does the mechanical barrier between basal and suprabasal layers help orient the division plane?

In biological systems, spindle orientation is intrinsically coupled to tissue mechanics. Cell shape, cortical tension, and intercellular forces influence spindle alignment and are themselves modified by the outcome of cell division [28, 3, 23, 29]. Moreover, external mechanical forces can directly reposition the mitotic spindle, highlighting that division orientation is mechanically sensitive rather than purely genetically prescribed [12, 44]. As a result, it is experimentally difficult to disentangle whether a given division orientation is the cause or the consequence of the surrounding mechanical environment.

Previous work has suggested the long axis of the cell might help orient cell divisions, and also that perpendicular divisions occur more frequently at early stages (E14), while planar divisions occur more frequently at later stages (E16) [22, 45, 3, 27, 29, 46]. To incorporate these experimentally observed trends into our analysis, we approximate the distribution of division orientations by grouping angles into three categories: planar (*θ* ∈ [0, *π/*6]), oblique (*θ* ∈ (*π/*6, *π/*3]), and perpendicular (*θ* ∈ (*π/*3, *π/*2]). Based on published data, we estimate the fractions of division orientations at early (*E*14) and later (≥ *E*15) developmental stages as

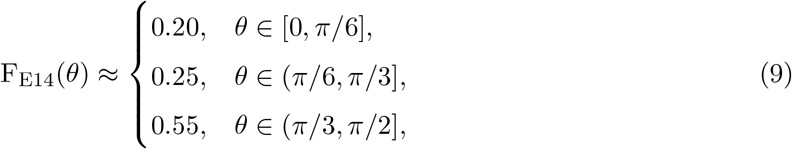

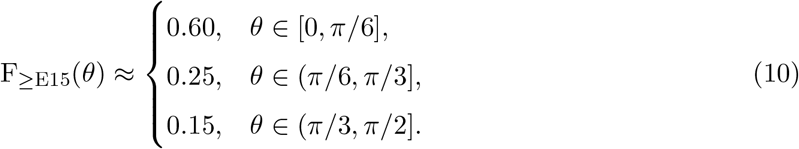

From these fractions, we construct normalized probability distributions for each developmental stage,

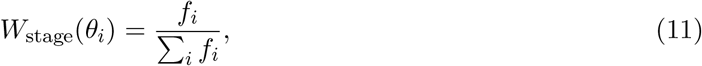

where *f*_*i*_ denotes the fraction of divisions within angular bin *θ*_*i*_. By construction, *W*_stage_(*θ*) satisfies ∑_*i*_ *W*_stage_(*θ*_*i*_) = 1.

Figure 2 (c) shows the normalized probability distribution of division angles for *E*14 and *E*16 estimated from the experimental data [45, 3, 27, 29, 46, 22].

**Figure 2:**
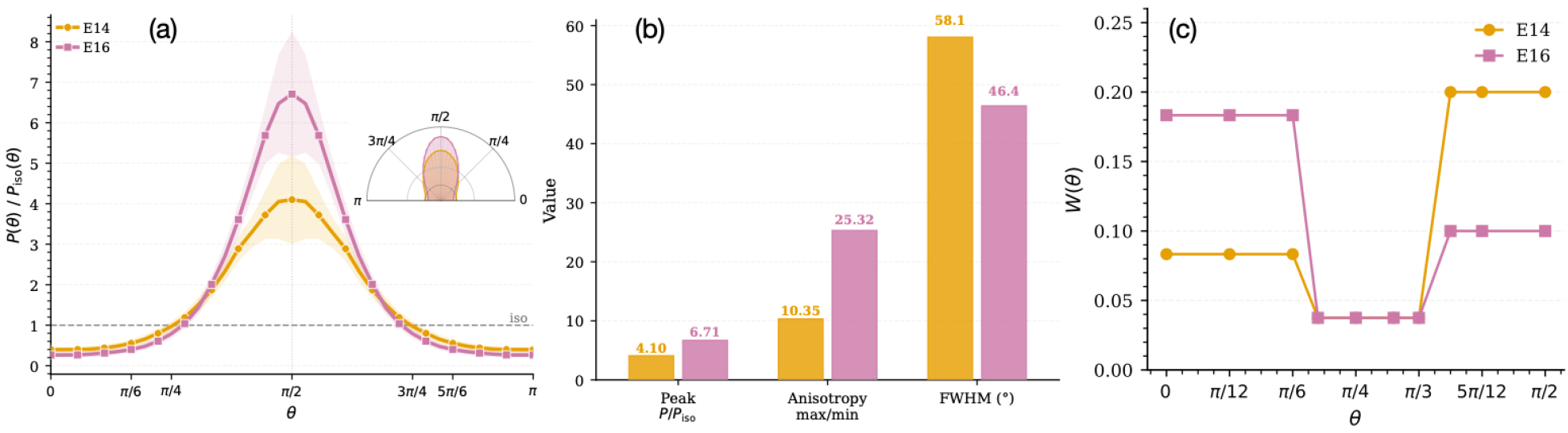
(a) Probability distribution of the orientation angle *θ* of the cell’s longest axis, obtained from an ellipsoidal fit and measured relative to the basal layer plane at developmental stages E14 and E16. Inset corresponds to a rose plot illustrating the angular distribution. Shaded regions represent the standard deviation from 50 independent simulation runs. (b) The peak probability, anisotropy ratio (max/min), and full width at half maximum (FWHM) for the distributions shown in panel (a) are indicated for developmental stages E14 and E16. (c) Normalized probability distributions of division angles for *E*14 and *E*16, obtained from experimental data [22, 45, 3, 27, 29, 46, 22].

We wonder if these changes in cell division orientation as a function of developmental stage could be a straightforward consequence of a changing long-axis orientation of growing cells constrained by a changing mechanical barrier. Previous studies [36, 45, 21] have shown that heterotypic interfacial tension can bias cell shape and thus the longest-axis orientation.

We therefore begin by examining the role of the mechanical energy barrier in setting the long-axis orientation of a growing cell. We randomly select a basal cell and grow it isotropically by increasing its target volume *V*_0_ according to

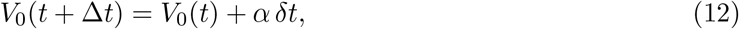

where *α* is the growth rate and *δt* = 0.01*τ* is the simulation time step. At any given time, only one basal cell is selected to grow, while all others have *α* = 0, ensuring stochastic selection in space and time.

To quantify how cell long-axis orientation depends on developmental stage, and thus on the mechanical barrier, we measure the probability distribution of the longest-axis orientation of the growing basal cell prior to division. Figure 2 (a) shows the distribution of the angle between the basal plane and the longest principal axis (obtained via ellipsoidal fitting) at stages *E*14 and *E*16. The distributions are defined on [0, *π/*2] and mirrored about *π/*2 to extend over [0, *π*], preserving the isotropic reference *P* (*θ*)*/P*_iso_(*θ*) = 1.

At both stages, cells preferentially align perpendicular to the basal layer (*θ* = *π/*2). However, the distribution at *E*16 is more sharply peaked, indicating stronger orientational order compared to *E*14.

This suggests that the mechanical environment increasingly constrains cells toward perpendicular orientations at later stages, likely due to enhanced heterotypic interfacial tension and basal-layer stiffness. In other words, as the skin epidermis matures, the enhanced interfacial coupling imposes stronger geometric and mechanical constraints on the basal cells, promoting the alignment of their longest axis relative to the basal plane. At *E*14, when heterotypic contacts are less pronounced, cells exhibit a wider range of orientations.

We further quantify this anisotropy using three metrics: (i) the peak normalized probability, (ii) the anisotropy ratio (maximum to minimum probability), and (iii) the full width at half maximum (FWHM). The peak probability increases from 4.10 (E14) to 6.71 (E16), while the anisotropy ratio rises from 10.35 to 25.32. Consistently, the angular spread decreases from 58.1° (0.32*π*) at E14 to 46.4° (0.26*π*) at E16, confirming stronger perpendicular alignment at later developmental stages.

In summary, we have shown that isotropic cell growth in the background mechanical environment favors *stronger* perpendicular long-axis orientation at E16 compared to E14. This *invalidates* our conjecture that the emerging mechanical barrier at E16 suppresses perpendicular cell divisions by lowering the probability that the long axis is oriented perpendicularly; the opposite effect is observed. Since perpendicular divisions are more common at E14, this suggests that there must be other mechanisms operating differentially between E14 and E16 to drive those perpendicular divisions. Additional regulatory mechanisms could include active spindle positioning, anisotropic cell growth, and/or molecular polarity cues.

### 3.4 Cell Division and Stratification

An advantage of our computational framework is that it allows us to impose division orientations and directly probe their impact on stratification dynamics and basal tissue mechanics. We prescribe division orientations independently of cell geometry and systematically quantify tissue growth and morphology as a function of the mechanical state. This controlled approach isolates the causal effect of division angle on stratification and tissue mechanics.

To make quantitative predictions about stratification dynamics as a function of division orientation and rate, we must first define when a cell becomes ‘stratified’. There are two types of stratification.

First, basal cells can spontaneously stratify due to shape and force fluctuations. To identify these events, at each time step, we compute the basal–suprabasal (BS) interface as the mean position of centroids of polygons shared between basal and suprabasal cells. A basal cell is considered to have stratified once its centroid lies above this interface and it no longer contacts the basement membrane, at which point its vertex model parameters are instantaneously altered to match those of suprabasal cells.

Cell divisions can either directly induce upward displacement, as a daughter cell is born with a centroid close to the BS interface and the mechanical environment drives it upward, or force fluctuations due to cell divisions can drive nearby cells to move upward. See SI Section 3 for a discussion of these two classes of events. We denote both of these types of events as spontaneous stratification of otherwise identical (homotypic) basal cells (orange cells in Fig. 3 (d-f)). We denote the number of such events as *N*_homo_.

**Figure 3:**
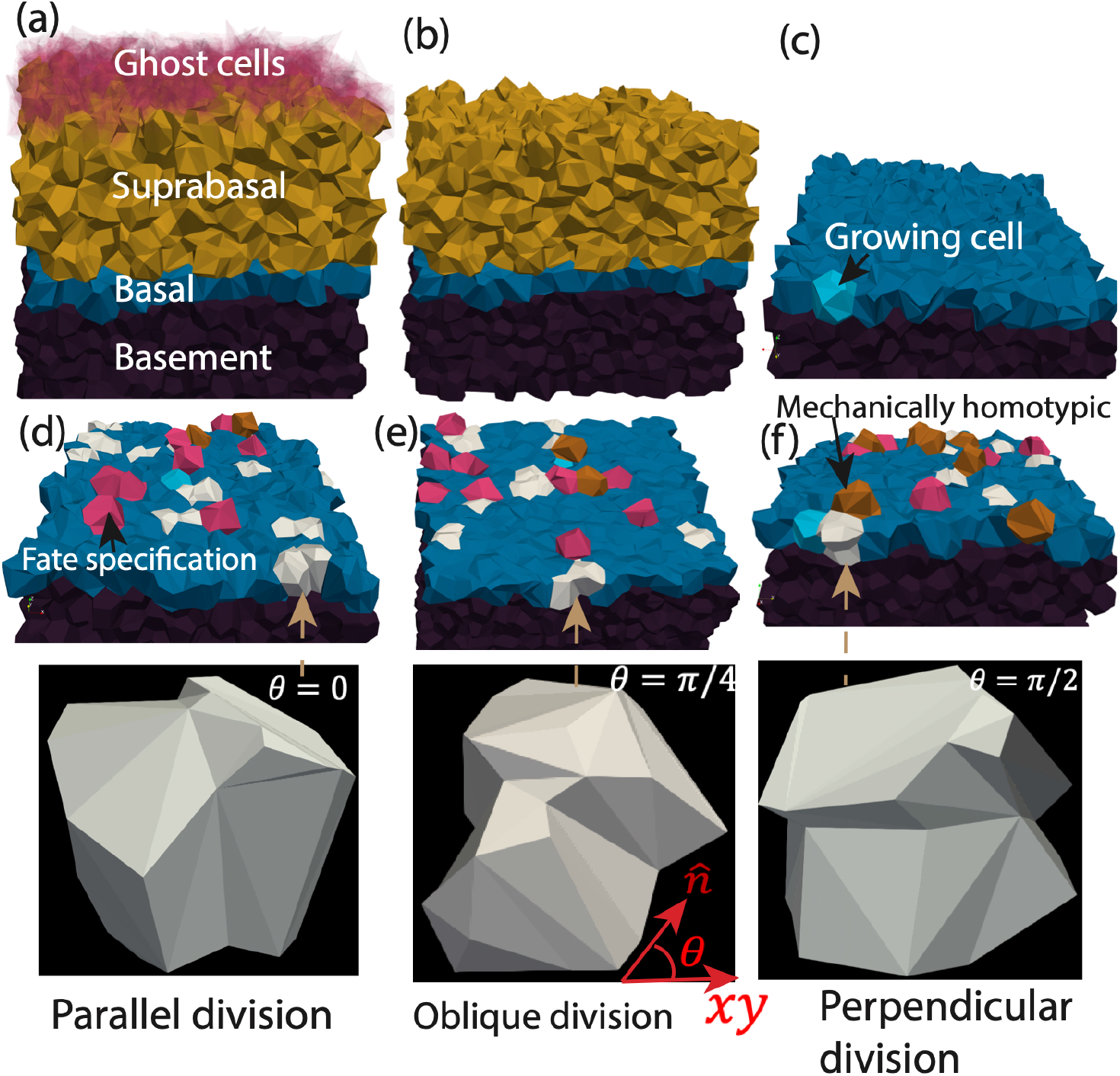
Visualization of the 3D vertex model of stratified epithelia illustrating stratification outcomes as a function of division orientation. (a) Snapshot of the full simulated tissue, showing the basement membrane (dark purple), basal layer (blue), suprabasal layer (gold), and ghost cells (transparent red). (b) Same snapshot as in (a) with ghost cells removed for clarity. (c) Proliferating basal cell (sky blue, black arrow) located in the basal layer. (d-f) Division orientation-dependent outcomes: parallel/symmetric (*θ* = 0, d), oblique (*θ* = *π/*4, e), and perpendicular (*θ* = *π/*2, f) divisions, respectively. White cells denote daughter cells after a cell division that remain in the basal layer; orange cells represent cells that have homotypically stratified, i.e. moved upward into the suprabasal layer due to the mechanical environment (see also Supplementary Movie M1); magenta-colored cells indicate cells that have been chosen to change their cell fate and mechanics so that they move upward, in order to maintain constant basal cell number density (see also Supplementary Movie M2).

Regardless of orientation angle, even with these spontaneous stratification mechanism, we find that repeated divisions lead to crowding and a strong overall increase in basal-layer density (see SI Section 2). This crowding leads to time-varying alterations to the mechanics of the basal layer and surrounding tissues as the simulation progresses. See SI Section 2 for an analysis of cell division in this “densifying” state.

Importantly, such overcrowding is generally not observed in wild-type embryos at later stages [3, 27, 29, 45, 46], and work by some of us has demonstrated that there is a notch-triggered cell fate specification pathway that leads to programmed delamination at later stages [45]. In addition, since our goal is to understand how cell divisions interact with tissue mechanics and geoemetry, it is important for the tissue to have a well-defined steady state mechanics and geometry during the simulations.

Therefore, we introduce a second mechanism for stratification that enforces density homeostasis in the basal layer and captures our observation of cell-fate-specification-driven delamination [45]. We term the protocol where we include this mechanism “homeostatic”, while the previous protocol (also described in SI Section 2) is termed a “densifying” protocol. In the homeostatic protocol, a basal cell is randomly selected to undergo a cell fate change and alter its mechanical properties to match those of a suprabasal cell. In agreement with previous work [45], we find that this switch in mechanical properties very robustly drives upward movement and delamination of the chosen cell (pink cells in Fig. 3 (d-f)). To ensure density homeostasis, at regular intervals *t*_*pop*_ the total number of basal cells *N*_basal_(*t*) is compared to its initial reference value *N*_basal_(0). Whenever the basal cell count has expanded by more than two cells, a random basal cell is converted to a suprabasal identity. The number of such events is denoted by *N*_fate_.

Together, *N*_homo_ and *N*_fate_ account for all stratification events, ensuring that the basal layer does not accumulate excess cells and remains in a homeostatic steady state, i.e. it maintains single-layer thickness. In practice, this means that the total number of stratification events, *N*_homo_ + *N*_fate_, is approximately equal to the number of cell divisions in the basal layer *N*_div_, and so we often normalize by *N*_div_ in order to compare systems at different division rates.

This strategy has no prescribed spatial bias; all basal cells have equal probability of being selected, reflecting the experimentally observed stochasticity of stratification events in squamous epithelia. Our homeostatic control scheme is intermediate between paired division–apoptosis approach [7] and with independent stochastic cell removal [43]. Cell divisions are not immediately compensated by imposed stratification; instead, excess basal cells are moved up only after the basal population exceeds a prescribed threshold, implementing delayed, collective regulation rather than instantaneous or independent turnover.

The stratified tissue architecture of the system is illustrated using representative snapshots, Fig 3(a-f), showing the spatial organization of its key components. Actively growing cells undergo division within basal layer, generating daughter cells that contribute to tissue stratification. To investigate the effect of division orientation, snapshots taken immediately before and after cell division are shown for three division angles, *θ* = 0, *θ* = *π/*4, and *θ* = *π/*2. All simulations are shown for a division rate *λ* = 0.49 divisions per basal cell per estimated day in the developmental stage E14.

Every stratification event induces an upward flux through the tissue. To maintain confluence and topological consistency without altering the mechanical state of the tissue, the top-most suprabasal cell is immediately converted into a ghost cell [47, 48]. Ghost cells are volume-less and surface-less placeholders that retain full topological connectivity (faces, edges, and vertices) but do not contribute to the mechanical energy or generate intercellular forces. Consequently, ghost cells do not exert elastic, contractile, or active forces on neighboring cells. Importantly, ghost cells remain eligible for T1 (reconnection) transitions, ensuring that local neighbor exchanges and surface remodeling proceed smoothly. This approach preserves global confluence, avoids artificial stress release at the tissue surface, and maintains a constant effective tissue height during stratification-driven flux. By pairing each basal-to-suprabasal transition with the removal of a suprabasal cell (conversion to ghost cell) at the apical surface, the tissue preserves a quasi-steady thickness and exhibits a well-defined upward flux of material.

Figure 4 (a) compares the fraction of homotypic stratification events (*N*_homo_*/N*_div_) for developmental stage *E*14 and *E*16. Our key observation is that *N*_homo_*/N*_div_ increases monotonically with *θ*, from symmetric divisions (*θ* ≈ 0) to asymmetric divisions (*θ* ≈ *π/*2). This trend is consistent across both developmental stages and indicates that division orientation is a primary determinant of stratification outcome.

**Figure 4:**
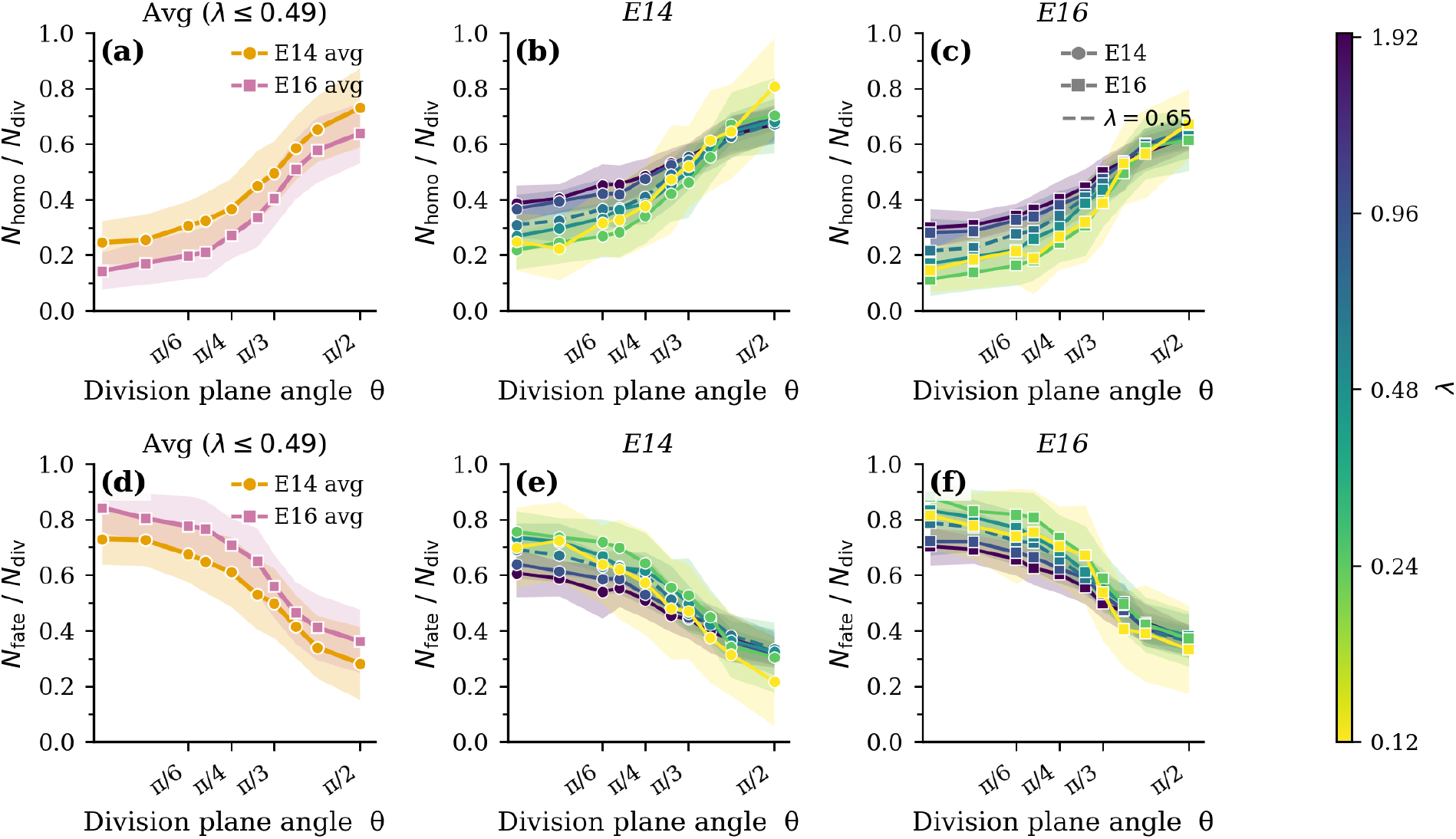
Stratification ratios as a function of division plane angle for varying division rates. (a) Comparison of the fraction of homotypic stratification events *N*_homo_*/N*_div_, averaged over quasi-static division rates *λ* ≤ 0.49 divisions per basal cell per estimated day, between developmental stages *E*14 and *E*16 across division plane angles. (b,c) *N*_homo_*/N*_div_ vs. division plane angle at stages *E*14 and *E*16, respectively, as a function division rate (purple = faster divisions, yellow = slower divisions). (d–f) Corresponding results for fraction of fate-based stratification events, *N*_fate_*/N*_homo_. In all panels, shaded regions represent the standard deviation from 40 independent simulation runs. See Supplementary Movies M3 and M4 for the dynamics of symmetric and asymmetric divisions, respectively.

This provides concrete confirmation from a well-vetted mechanical model for an idea that is intuitively obvious and already posited in experimental manuscripts: low-angle (planar) divisions tend to expand or maintain the basal layer, contributing weakly to stratification, whereas high-angle (oblique to perpendicular) divisions preferentially displace one daughter cell upward away from the basal layer, increasing suprabasal cell production and thus promoting stratification. The gradual rise in *N*_homo_*/N*_div_ with *θ* indicates a continuous, rather than binary, relationship between division orientation and stratification outcome, suggesting that intermediate angles contribute partially to stratification rather than enforcing a strict planar versus-perpendicular distinction.

Although we group all homotypic stratification events together, the time required for stratification varies systematically with orientation angle, with parallel divisions requiring more time to stratify. See SI Section 3 for more details.

Across all division angles, division yields greater stratification at *E*14 than at *E*16, a trend consistent with the stage-dependent mechanical landscape of the tissue. At *E*14, the effective mechanical barrier at the basal–suprabasal interface is weak, characterized by lower stiffness of the basal layer, and reduced heterotypic interfacial tensions (*σ*_*a*_). In this regime, daughter cells produced by oblique, perpendicular, or even planar divisions can more readily rearrange, undergo T1 transitions, and cross the interface, so a larger fraction of divisions is successfully converted into stratified outcomes. In contrast, at *E*16 the basal layer has stiffened and together with the increased heterotypic tension generates a stronger mechanical barrier. This elevated barrier increases the energetic cost of upward displacement and interfacial deformation, thus suppressing delamination and reducing the stratification yield per division even when the division angle is favorable.

This process also depends slightly on the rate of cell divisions. Figure 4 (b,c) show that for timescales shorter than 200 natural time units (∼ 20 minutes), faster cell divisions allow more spontaneous homotypic stratification events when angles are close to parallel. The behavior is quite similar between simulations when the cell division rate is smaller than 0.49, suggesting that cell divisions are slow enough that they do not induce fluctuations above the baseline fluctuations *D*_*t*_ (Eq. 2).

Since we have implemented a protocol to maintain density homeostasis, *N*_fate_ must necessarily follow precisely the opposite trend as *N*_homo_ (Fig 4 (d-f); our model predicts that cell fate specification is required more frequently at E16 compared to E14 to maintain homeostasis at a specified cell division angle. Moreover, cell fate specification is required more frequently when cell divisions are oriented parallel to the basement membrane compared to perpendicularly. Both predictions are consistent with experimental observations [45].

An obvious question is whether our protocol implementing density homeostasis strongly affects these results. SI section 2 compares results for *N*_homo_ and *N*_fate_ between this homeostatic protocol and the “densifying” protocol where there is no cell-fate-driven delamination. We compare the two at *λ* = 0.49 divisions per basal cell per estimated day (SI Fig S1(a,b)), which is close to the rate of 0.7 divisions per basal cell per day reported for mouse epithelia between E14 and E16. SI Fig S1(c) shows that *N*_homo_ is remarkably consistent between the homeostatic and densifying protocols at E14, and still fairly similar across all orientation angles even at E16. This suggests that the moderate densification consistent with experimental observations is not sufficient to shift material properties or alter homotypic stratification mechanisms.

Taken together, these results indicate that division orientation alone does not fully determine the efficiency of stratification; instead, the developmental mechanical context modulates how angled divisions translate into stratified outcomes. Our data are consistent with the hypothesis that the emergence of a stronger mechanical barrier at later developmental stages suppresses homotypic stratification relative to the early, more permissive *E*14 state [45].Furthermore, faster cell divisions (occuring over timescales shorter than every ∼ 20 minutes) can enhance the effective mechanical fluctuations and promote spontaneously delamination when the division angle is close to parallel.

### 3.5 How do cell division orientation and rate alter basal layer fluidization and mechanics?

The results of the previous section already suggest that faster cell divisions may alter mechanical properties of the basal layer. This is consistent with previous studies on 2D tissue simulations [25, 7], which reported cell-division induced fluidization.

To quantify this effect, we measure the relative squared separation Δ*d*^2^(*t*) (see Section 2.2) between daughter cells that remain in the basal layer as a function of time, as shown in Fig. 5 (a,b). To characterize these dynamics, we analyze the time evolution of Δ*d*^2^(*t*) (Fig. 5, (a,b)). At long times and faster division (*λ* = 1.95 divisions per basal cell per estimated day), we observe diffusive behavior,

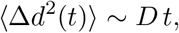

from which we extract an effective diffusion coefficient *D* via linear fits to the blue shaded regions in Fig. 5, (a,b)).

**Figure 5:**
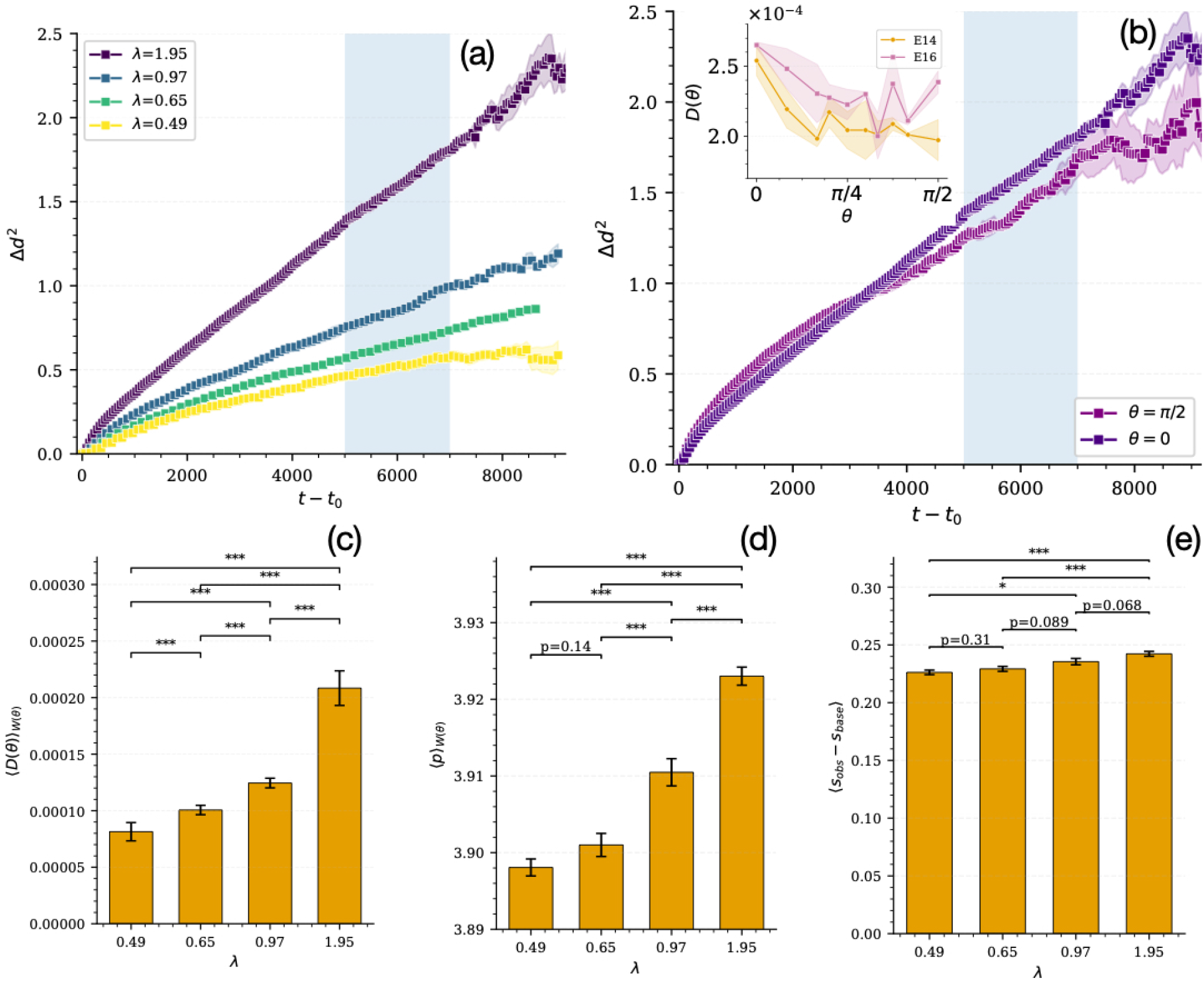
Faster divisions and planar-orientated divisions drive fluidization of the basal layer. (a) Time evolution of the squared separation Δ*d*^2^(*t*), defined in Section 2.2, between daughter cells that remain in the basal layer for varying division rates, as a function of the time since division *t* − *t*_0_. (b) Squared separation for planar (*θ* = 0) and perpendicular (*θ* = *π/*2) divisions. Inset: diffusion coefficient as a function of division angle *θ* at developmental stages E14 and E16 for *λ* = 1.95 divisions per basal cell per estimated day. The methods for computing error bars for these panels are described in the main text. (c-e) Simulation data for parameters corresponding to E14 weighted by distribution of cell orientations experimentally observed in E14 embryos (*W* (*θ*)) from Fig. 2(c). (c) Orientation-weighted diffusion coefficient using experimentally measured division-angle distributions. (d,e) Orientation-weighted average 2D and 3D basal cell shapes, respectively. The error bars in (c-e) are the standard deviation of the mean over the orientation angles.

We estimated the error on the measured squared separation by binning the squared separation Δ*d*^2^ between daughter cells that remain within the basal layer into time intervals of width Δ*t*. In each bin, the mean is computed over all samples, while the uncertainty (standard error) is estimated by bootstrap resampling and subdividing the full dataset (*N* = 40 samples) into *n* = 5 independent subgroups (8 samples each). The binned Δ*d*^2^ is computed for each subgroup, and the error is taken as the standard deviation of the subgroup means divided by squared root of the number of subsampled subgroups.

The diffusion coefficient is extracted from a linear fit of the binned MSD within the prescribed long time diffusion window (shaded in blue in Fig 5(a,b)). Its uncertainty is obtained by computing the diffusion coefficient for each subgroup and reporting the standard deviation of the subgroup estimates.

We find that the dynamics depend on both division rate and orientation. At low division rates, Δ*d*(*t*) exhibits subdiffusive behavior, indicating that daughter cells remain constrained in a solid-like environment, similar to the solid-like behavior reported in 2D in Ref. [7]. This is also consistent with recent work in models for 3D layered tissues [39, 16], where it is reported that the tissue remains solid-like in the absence of sufficiently large division rates.

However, we find that further increasing the division rate enhances fluctuations and local rearrangements, and for sufficiently fast divisions (*λ* = 1.95 divisions per basal cell per estimated day), the dynamics become diffusive, signaling fluid-like behavior of the basal layer. Parallel divisions induce more diffusion in the basal layer than perpendicular divisions (Fig. 5, (b)).

These results indicate that division orientation controls how mechanical stress is injected into the basal tissue. Planar divisions generate in-plane displacements that directly perturb local packing, enhancing fluctuations in the basal layer and promoting diffusive dynamics. In contrast, perpendicular divisions primarily displace material into suprabasal layers, relieving basal stress and producing fewer in-plane rearrangements. Thus, planar divisions act as an stronger source of active mechanical fluctuations in the basal layer.

This behavior is captured by the diffusion coefficient *D*, which increases significantly with division rate (SI Fig. S3(a)), consistent with division acting as a source of active noise that drives fluidization. A systematic analysis of diffusion as a function of orientation angle (inset of Fig. 5(b)) confirms that *D*(*θ*) is highest for parallel division and decreases nearly monotonically with increasing cell division angle. Perhaps counterintuitively, *D*(*θ*) is higher across all division angles in the E16 background compared to E14. We conjecture that the stronger mechanical barrier at E16 may restrict cell-division-induced fluctuations to the basal layer, thereby enhancing them compared to E14. Additional data for the dependence of *D* on cell division rate, orientation angle, and developmental stage is shown SI Fig. S3(a).

Although we have plotted data as a function of orientation angle and developmental stage independently, experiments have demonstrated that the cell division angle changes systematically with developmental stage (Fig 2 (c)). Therefore, to develop an experimentally relevant prediction based on our simulations, we weight the simulation data for the diffusion at each angle by its observation probability in experiment at that stage of development:

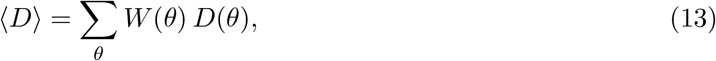

where *W* (*θ*) is the measured distribution of division angles for that stage (see Fig 2 (c)). This reveals a clear increase in fluidization with division rate. The data for ⟨*D*⟩ as a function of division rate in the E14 mechanical background is shown in Fig. 5(c).

In addition, we compute and analyze both 2D and 3D basal-cell shapes (see SI Section 4 for definitions), which have been shown to correlate with fluidity in 2D [32, 2] and 3D [26, 48] to further characterize the coupling between division geometry and basal-layer fluidization.

We perform a similar orientation-weighted average of the 2D and 3D cell shapes at E14. We find the diffusion constant is well-correlated with 2D basal-layer shape measurements (Fig. 5, (d)), which exhibit increased cell shape at higher division rates. Interestingly, in these layered 3D systems, the observed 2D shape index is more strongly correlated with the diffusion constant than the 3D shape index (Fig. 5(e)), which has been shown to govern diffusion in disordered 3D tissues [26, 48].

As shown in SI Fig S3(c,d), the higher fluidity at E16 compared to E14 also shows up in the orientation-weighted average of cell shapes and diffusion constants. Although the basal layer is intrinsically stiffer at E16 (higher Δ*s*), our model predicts that in-plane fluidization is enhanced by fast cell divisions, due to both the stronger impact of fluctuations and the higher prevalence of planar divisions.

Together, these findings demonstrate that cell division acts as a source of active mechanical fluctuations in the basal layer that can fluidize an otherwise solid-like tissue, and that this fluidization is enhanced by planar-oriented divisions and the mechanical energy barrier that emerges at E16. Moreover, 2D shape index (measured in the plane parallel to the basement membrane) provides a robust readout of this fludization.

## 4 Conclusion and Discussion

In this work, we extend a recently developed three-dimensional (3D) vertex model of stratified epithelia to incorporate basal stem-cell divisions, enabling us to investigate the mechanical feedback between the basal-suprabasal mechanical barrier, division plane orientation, and epidermal multilayering. By systematically varying division orientation and rate, we establish a quantitative framework linking division geometry to stratification dynamics, basal-layer fluidization, and tissue organization across developmental stages.

Because epithelial remodeling involves coordinated rearrangements along the apical–basal axis that are not captured by purely two-dimensional descriptions [20], our 3D framework is essential to resolve how local cell-level processes give rise to tissue-scale behavior. By integrating our model with experimental measurements, including droplet-based mechanical calibration [40] and tissue-level observations [45], we establish a direct correspondence between simulation parameters and experimentally accessible quantities, enabling quantitative comparison between theory and experiment.

Our initial hypothesis was that the stage-dependent mechanical barrier at the basal–suprabasal interface, transitioning from a compliant, fluid-like state at early developmental stages (*E*14) to a stiffer, more resistive barrier at later stages (*E*15–*E*16), might regulate division orientation. We postulated that since heterotypic interfaces have been shown to alter cell shapes and orientations, the emerging mechanical energy barrier could change the orientation of the long axis of the cell and thereby mechanically regulate the axis of division. However, we find the opposite: isotropically growing cells orient perpendicularly *more often* at E16 compared to E14. Our results demonstrate that barrier-driven long-axis orientation is *not* responsible for enhanced perpendicular divisions observed at E14. Instead, division orientation must be regulated by additional stage-specific cues, likely involving biochemical signaling, polarity pathways, and active spindle positioning mechanisms. Therefore, an important next step would be to incorporate explicit models of spindle positioning and polarity signaling, such as planar cell polarity pathways, cortical force generators, and cytoskeletal asymmetries, into the 3D vertex framework. This will enable us to make quantitative predictions of the most important molecular and biophysical mechanisms (forces) that control division orientation in a stage-dependent manner and vice-versa (make quantitative predictions of the feedback-loop). We next confirm the intuitive hypothesis that division orientation provides a direct control over stratification: planar divisions predominantly maintain the basal layer, whereas oblique and perpendicular divisions promote upward displacement of daughter cells and enhance stratification. This effect is more pronounced at earlier developmental stages, where the mechanical barrier is weaker and cells are more permissive to rearrangements. These findings indicate that while the mechanical environment defines a permissive landscape, division orientation acts as an instructive cue that determines tissue architecture.

Beyond stratification, we show that cell division acts as a source of active mechanical fluctuations that regulate basal-layer fluidization. Increasing division rates enhance cell rearrangements and drive a transition toward fluid-like behavior, consistent with concepts from active matter and glassy systems [7, 25]. Importantly, the efficiency of this fluidization depends on division orientation: planar divisions inject mechanical activity directly within the basal plane, whereas perpendicular divisions redistribute stresses toward suprabasal layers. Thus, division geometry not only controls stratification but also modulates the effective mechanical behavior of the tissue.

Our findings highlight that division orientation cannot be understood solely from passive mechanical cues, but instead arises from the coupling between mechanics and active cellular regulation.

Numerous experimental and theoretical studies have identified tissue jamming/rigidity and interfacial tension as key regulators of cancer progression and metastasis [35, 13, 32, 17]. However, the mechanisms and timing of carcinoma initiation within the basal layer of the epidermis remain poorly understood. Therefore, an important application of this framework would be in understanding pathological conditions, particularly cancer initiation and progression. Misoriented cell divisions are known to disrupt tissue architecture and can contribute to tumor initiation and progression. Extending our model to include aberrant polarity signaling, altered interfacial tensions, or dysregulated proliferation will allow us to investigate how defects in division orientation lead to loss of tissue organization, enhanced fluidization, and invasive behavior. In particular, coupling division orientation with changes in mechanical heterogeneity and active forces may provide insight into how transformed cells escape normal stratification constraints and invade surrounding tissues. More broadly, our work establishes a quantitative and predictive framework for studying how cell-level processes – such as division, mechanics, and polarity – integrate to control tissue morphogenesis.

By bridging molecular regulation, cellular mechanics, and tissue-scale dynamics, this approach opens new avenues for understanding developmental processes and their failure in disease.

## Author Contributions

**Somiealo Azote Epse Hassikpezi**: Conceptualization, Data curation, Formal analysis, Investigation, Methodology, Software, Validation, Visualization, Writing–original draft, Writing–review & editing. **Rajendra Singh Negi**: Conceptualization, Data curation, Formal analysis, Investigation, Methodology, Software, Validation, Visualization, Writing–original draft, Writing–review & editing. **No (Arnold) Chen**: Methodology, Writing–review & editing. **M. Lisa Manning**: Conceptualization, Funding acquisition, Investigation, Methodology, Project administration, Resources, Software, Supervision, Validation, Writing–original draft, Writing–review & editing.

## Acknowledgments

We would like to thank Sara Wickstrom, Carien Niessen, and their respective laboratories, for conversations and feedback on our modeling approaches to help them capture features from experimental observations.

## Supplemental Data

Supplementary Material should be uploaded separately on submission, if there are Supplementary Figures, please include the caption in the same file as the figure. LaTeX Supplementary Material templates can be found in the Frontiers LaTeX folder.

## Data Availability Statement

The datasets generated and analyzed for this study can be found in the associated dryad data repository [LINK will be available upon acceptance of manuscript].

