## Supplemental Text and Figures for "A Predictive Model for Coupling Cell Division Orientation to Tissue Mechanics During Epithelial Morphogenesis"

April 17, 2026

#### 1 Model parameters for dynamic 3D vertex model

As discussed in the methods section in the main text, the data reported here are from a 3D layered vertex model with active force fluctuations and overdamped dynamics. There are multiple vertex model parameters, and here we fix the values of those parameter to match data from experiments on stratified epithelia in the developing mouse embryo, using a process described in detail in that previous work [3]. In particular, some of us previously found that there are two key parameters that differ substantially between stages E14 and E16: heterotypic interfacial tension at the apical side of basal cells $\sigma_a$ and basal cell stiffness, $\Delta s = 5.4 - s_{basal}$. There is also a small difference in the wetting tension with the basement membrane, $\sigma_b$. Other vertex model parameters are identical between the two simulations, and fixed at values used previously [3]. For completeness, we list the vertex model parameters used in simulations for this manuscript in Table S1 below:

#### 2 Densifying protocol: Cell division without cell fate specification

In the main text, we focused on a "homeostatic" protocol for simulations that were maintained in steady state, with constant cell number density, via cell fate specification. We made this choice to allow us to study the impact of oriented cell divisions in a tissue with fixed background properties.

However, an alternate approach is to allow cell divisions to occur without any explicit cell fate decisions/forced delaminations, and in general this will lead to a time-varying increase in the number of basal cells. We term this protocol a "densifying" protocol. As has been reported previously [1], such crowding is expected to eventually drive mechanical instabilities that can lead to enhanced homotypic stratifications (stratification without the need to change cell-cell interactions/mechanics), that could thereby suppress further density increases. Although previous work by some of us suggests this is not the case at later stages (E16.5) [3], it could be a relevant mechanism at earlier stages (E14.5).

Table 1: Simulation Parameters for E14 and E16

| **Parameter** | **E14** | **E16** |
|---|---|---|
| $\mu$ | 1.0 | 1.0 |
| $k_v$ | 10.0 | 10.0 |
| $k_s$ | 1.0 | 1.0 |
| $D_t$ | 1.0 | 1.0 |
| $s_{basement}$ | 5.14 | 5.14 |
| $s_{basal}$ | 5.40 | 5.26 |
| $s_{suprbasal}$ | 5.77 | 5.77 |
| $v_{basement}$ | 1.0 | 1.0 |
| $v_{basal}$ | 1.0 | 1.0 |
| $v_{suprbasal}$ | 1.0 | 1.0 |
| $\theta$ | $0 - \pi/2$ | $0 - \pi/2$ |
| $T$ | 0.002 | 0.002 |
| $T_{\text{bot}}$ | 0.002 | 0.002 |
| $\sigma_b$ | 0.062 | 0.067 |
| $\sigma_a$ | 0.044 | 0.116 |
| $\lambda$( divisions per basal cell per estimated day) | $0.12 - 1.95$ | $0.12 - 1.95$ |

To investigate this, we implement a model of cell division that excludes explicit cell fate specification during stratification. Here we focus on simulations with a division timescale of $t_{div} = 200$, as it is close to the rate reported in the literature for basal cells in E14-E16 (about 0.7 divisions per basal cell per day), see main text Section 3.1. We study this process for $10,000\tau$ simulation time units, which is about 17 hours and similar to an embryonic day (i.e. the amount of time between stage E14 and E15).

In this framework, the density of the basal layer is allowed to vary and stratification arises solely through homotypic interactions, without any induced changes in cell fate to facilitate layering. Here, the density of the basal layer is governed by the division angle, $\theta$. Perpendicular divisions promote greater homotypic stratification, whereas in-plane divisions result in reduced homotypic stratification, leading to an increase in basal layer cell density. Figure 1 shows the evolution of basal cell numbers as a function of time $\tau$ for both protocols. The left panel (a) corresponds to the "homeostatic" constant-density case, where the basal population $N_{\text{basal}}$ remains approximately constant for both division angles. In contrast, the right panel (b) shows the "densifying" protocol, where the number of basal cells increases over time and the fate specific stratification is not applied randomly. Perpendicular divisions without fate change lead to a slow increase in density, as the added basal cells are largely balanced by stratification. By contrast, parallel (in-plane) divisions result in a continuous increase in basal cell density.

Figure 1(c) shows that, perhaps surprisingly, densification appears to have little impact on stratification at $E14$. In other words, the fraction of cells that stratify due to homotypic interactions in these "densifying" simulations is nearly the same as in the "homeostatic" simulations where we

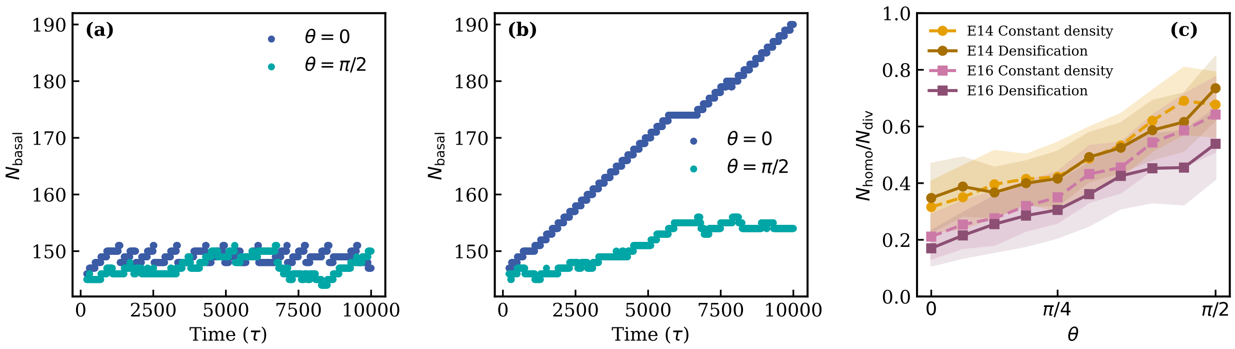


Figure 1: (a,b) Basal cell counts $N_{\text{basal}}$ as a function of time in natural units $\tau$ for "homeostatic" constant density (a) and "densifying" (b) protocols at developmental stage $E14$ with a division rate of $\lambda = 0.49$ divisions per basal cell per estimated day. Dark blue denotes parallel divisions ($\theta = 0$) and light blue denotes perpendicular divisions ($\theta = \pi/2$). Both panels share the same vertical scale to enable direct comparison of the growth dynamics between protocols. (c) Homotypic stratification fraction $N_{\text{homo}}/N_{\text{div}}$ as function of division angle for developmental stage $E14$ and $E16$, comparing homeostatic(constant density) vs. densifying protocols. For densifying protocols, the stratification is measured between $t = 5000 - 10,000\tau$, i.e when the densification is well underway. Shaded regions represent the standard deviation from 40 independent simulation runs.

forced cell delaminations; in the "densifying" case there are simply more cells that stay in the basal layer instead of being forced to delaminate.

For our simulations in the E14 background, there is orientation-dependent crowding induced at the end of $10,000$ natural time units $\sim 17$ hours, which is driven by $N_{div} = 49$ total divisions. Fig 1(b) shows that in that time window, the densifying protocol generates an extra 43 cells ($\approx 29\%$ more ) in the basal layer for parallel divisions with $\theta = 0$, and 7 additional cells ($\approx 5\%$ more ) in the basal layer for perpendicular divisions $\theta = \pi/2$. However, as shown by the orange E14 curves in Fig 1(c), these crowding effects are not sufficient to induce extra homotypic stratification.

For the $E16$ background mechanics, the greater densities generated in the "densifying" simulations slightly decrease the number of homotypic stratification events for all values of division orientation, although the effect is fairly small.

Taken together, this suggests that in both the E14 and E16 mechanical backgrounds, moderate increases in cell density (of up to about 30%) do not substantially change the stratification results compared to the "homeostatic" protocol reported in the main text at constant cell density. For this set of vertex model parameters, this means that a moderate change in density does not induce an instability that enhances homotypic stratification. Of course, if we were to increase the rate of cell division further (i.e. $\lambda > 0.49$ divisions per basal cell per estimated day), densities would increase further over the same time window and could eventually lead to mechanical instabilities [1]. This would be an interesting avenue for future research.

#### 3 Delay time for homotypic stratification

As described in the main text, homotypic stratification events were identified from cell-state time series extracted from each simulation run. A homotypic stratification event is defined as a transition in which a basal cell with identical properties to its neighbors moves upward into the suprabasal layer due to mechanical interactions. For simplicity, in the main text we focused on the total number of homotypic events that occurred at any time after a cell division, $N_{homo}$, but there is additional structure in how these events occur as a function of time.

For each homotypic stratification event, the delay time $\Delta t_d$ was defined as the interval between the most recent time the cell entered the daughter state (after divsion event) and the time at which the cell centroid breached the suprabasal layer. Two mechanistically distinct classes were distinguished: (i) immediate stratifiers, in which the cell transitioned directly, corresponding to events with $\Delta t_d \leq 10\tau$ to account for finite temporal resolution in the simulation output, and (ii) delayed stratifiers, in which the cell resided in the basal layer for a measurable duration prior to stratification, yielding $\Delta t_d > 0$.

###### 3.0.1 Zero-Inflated Gamma Model

The distribution of delay times across all mechanical stratification events was modeled using a Zero-Inflated Gamma (ZIG) distribution. This model was chosen because the data contain a discrete point mass at zero (immediate stratifiers) superimposed on a continuous, strictly positive, right-skewed distribution (delayed stratifiers), a structure that cannot be captured by a standard unimodal distribution. The ZIG model is defined as a two-component mixture [2]:

$$P(\Delta t) = p_0 \delta(\Delta t = 0) + (1 - p_0)\Gamma(\Delta t | k, m), \Delta t > 0 \tag{1}$$

where $p_0 \in [0, 1]$ is the weight representing the probability of immediate stratification, $\delta$ is the delta function, and $\Gamma(k, m)$ is the two parameter Gamma distribution with shape parameter $k > 0$ and scale parameter $m > 0$. The mean and variance of the non-zero component are $\mu_{nz} = km$ and $\sigma^2_{nz} = km^2$ respectively.

Figure 2 shows the dependence of the fitted ZIG parameters on the division angle for two developmental stages, $E14$ and $E16$. Figure 2 (A) shows the zero-inflation parameter $p_0$, which represents the probability of immediate stratification. In both developmental stages, $p_0$ varies systematically with $\theta$, indicating that the likelihood of immediate stratification depends on the angle of division. For symmetric divisions ($\theta \approx 0$), the cleavage plane produces two daughter cells that are mechanically and geometrically equivalent with respect to their surrounding environment. This symmetry suppresses any immediate directional bias or force imbalance that could drive stratification at the moment of division. As a result, stratification usually does not occur instantaneously, leading to a reduced probability of immediate stratification (lower $p_0$) and longer delay times. In this regime, additional time is required for stochastic fluctuations, local rearrangements, or external mechanical cues to break the initial symmetry before stratification can proceed. In contrast, highly

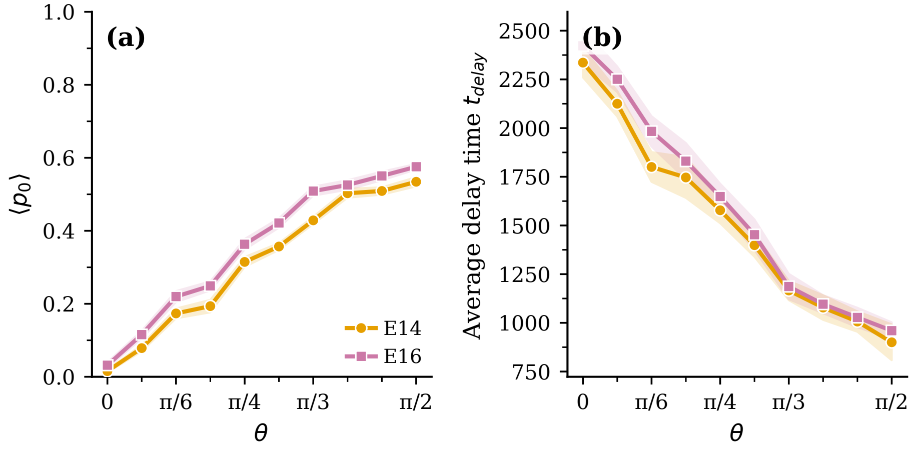


Figure 2: **Dependence of stratification delay statistics on the division angle $\theta$ at a fixed division rate.** (a) Zero–inflation parameter $p_0$, representing the probability of immediate stratification ($\Delta t \leq 10\tau$), as a function of the division angle $\theta$ for developmental stages E14 (orange) and E16 (pink) at $\lambda = 0.49$ divisions per basal cell per estimated day. (b) Mean delay time of the nonzero component of the distribution as a function of $\theta$ for the same developmental stages and rate. Shaded regions denote the standard error of the mean (SEM). Only selected values of $\theta$ are labeled on the horizontal axis for clarity.

oblique divisions ($\theta \approx \pi/2$) inherently introduce strong geometric and mechanical asymmetries between daughter cells. This built-in asymmetry increases the likelihood of immediate stratification, reflected by a higher $p_0$, and reduces the characteristic delay time for the remaining delayed events. Intermediate division angles interpolate between these two limits, generating a mixture of immediate and delayed stratification events. The smooth angular dependence of both $p_0$ and the mean nonzero delay time indicates that the division angle acts as a continuous control parameter that regulates how efficiently mechanical asymmetry is generated at division.

Figure 2 (B) shows the mean delay time of the nonzero component of the distribution, $\mu_{nz}$, as a function of $\theta$. This quantity characterizes the typical waiting time for delayed stratification events. For symmetric divisions ($\theta \approx 0$), the two daughter cells are created in a mechanically equivalent configuration. In the absence of an intrinsic asymmetry at division, stratification cannot proceed immediately and instead requires a subsequent symmetry-breaking process. This may arise from stochastic fluctuations, gradual force redistribution, or interactions with neighboring cells. As a consequence, stratification occurs only after a prolonged waiting period, leading to larger mean delay times for symmetric divisions. In contrast, highly oblique divisions ($\theta \approx \pi/2$) generate a pronounced mechanical asymmetry at the moment of division. Differences in geometry, force transmission, or coupling to the surrounding tissue provide an immediate directional bias that facilitates stratification. When stratification is delayed in this regime, the required mechanical reorganization is minimal, resulting in significantly shorter delay times compared to symmetric divisions. Intermediate angles exhibit delay times that interpolate between these two limits, consistent with a gradual increase in division-induced asymmetry as $\theta$ increases.

#### 4 Division-induced basal-layer fluidization

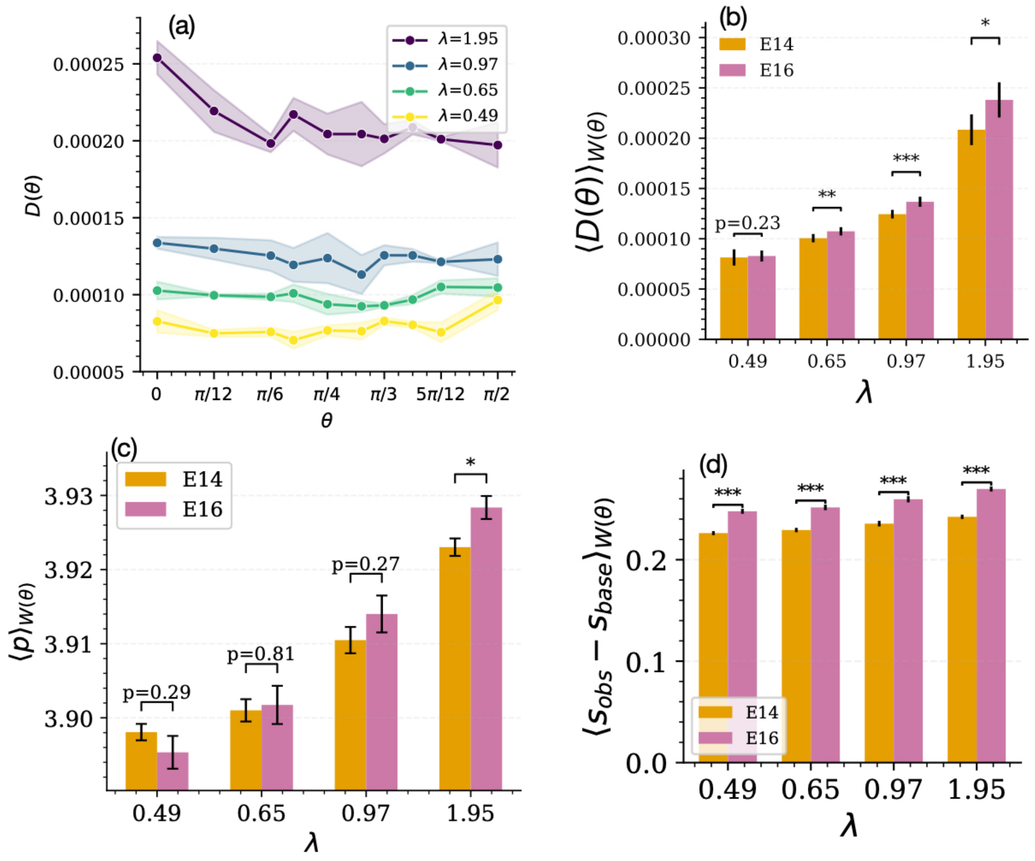


Figure 3: (a) Diffusion coefficient from daughter-cell separation as a function of division rate and angle $\theta$ at $E14$. (b–d) Probability-weighted averages (using experimental orientation distributions) of diffusion coefficient (b), 2D shape index (c), and change in observed 3D shape index or stiffness (d) at $E14$ and $E16$ across division rates. Error bars denote standard deviation over orientations.

We compute the squared separation between daughter cells (Equation 6 in the main text) within the basal layer as a function of time, division orientation, and division rate, and use this to estimate effective diffusion (rearrangement) rates. Figure 3(a) shows the diffusion coefficient extracted from the MSD for different division rates as a function of the division angle $\theta$. The diffusion depends strongly on division rate, with faster divisions leading to increased rearrangements. In contrast, the dependence on division orientation is weak and remains nearly constant.

The orange bars in Fig 3(b) show the diffusion data for E14 from Fig 3(a), averaged over orientation angle. Specifically, the average is weighted by the probability of observing a given orientation at E14 (see main text Eq. 13 and Fig 2(c).) The pink bars show the same averaging for E16 simulation data. As already discussed in the main text for E14 data, the diffusion is larger when there is faster rates of division (i.e. $\lambda = 1.95$ divisions per basal cell per estimated day). The E16 data shows the same overall trends, and for the faster division rates E16 is slightly more diffusive than E14.

We next examine whether cell shape correlates with basal-layer rearrangement dynamics. To this end, we quantify changes in basal-layer stiffness via $\Delta s = s_{\rm obs} - s_{\rm base}$, where $s_{\rm obs}$ is the 3D shape of basal cells in the presence of divisions and $s_{\rm base}$ is the corresponding shape in the absence of divisions. The 3D shape index is defined as $s = A/V^{2/3}$, where $A$ and $V$ are the cell surface area and volume, respectively. The 2D shape index is defined as $p = P/\sqrt{A_p}$, where $A_p$ and $P$ are the area and perimeter of the mid-plane cross-section of a basal cell.

Figure 3(c,d) compares 2D and 3D shape indices at developmental stages $E14$ and $E16$, again weighted by then probability of an orientation being experimentally observed at each stage, across division rates. Both metrics show increased fluidization with increasing division rate, as reflected by higher shape indices, although the 2D metric is much more strongly correlated with diffusion than the 3D metric. This trend is consistent across stages, with a slight enhancement of fluidization and higher shape index at $E16$ compared to $E14$.

### 5 Description of Supplementary Movies

- M1: Homotypic stratification of basal cells at developmental stage $E14$ and division rate of $\lambda = 0.65$ divisions per basal cell per estimated day. Orange-colored cells represent cells that have undergone homotypic stratification, acquiring a differentiated suprabasal type.

- M2: Stratification based on basal cell fate specification at developmental stage $E14$ and $\lambda = 0.65$ divisions per basal cell per estimated day. Magenta-colored cells indicate cells that are mechanically induced to stratify in order to maintain a constant basal layer density.

- M3: Dynamics of symmetric division ($\theta = 0$) at developmental stage $E14$ with $\lambda = 0.65$ divisions per basal cell per estimated day.

- M4: Dynamics of asymmetric division ($\theta = \pi/2$) at developmental stage $E14$ with $\lambda = 0.65$ divisions per basal cell per estimated day
